# Electrical potential spiking of kombucha zoogleal mats

**DOI:** 10.1101/2022.08.03.502684

**Authors:** Andrew Adamatzky

## Abstract

A kombucha is a sugared tea fermented by a symbiotic community of over twenty species of bacteria and yeasts. The community produces and inhabits cellulosic gelatinous zoogleal mats. We studied electrical activity of the kombucha mats using pairs of differential electrodes. We discovered that the mats produce action like spikes of electrical potential. The spikes are often grouped in the trains of spikes. Characteristics of the spikes and trains of spikes are presented. We demonstrated that electrical responses of kombucha mats to chemical, electrical and optical stimulation are distinctive and therefore the mats can be used as sensors, or even unconventional computing devices.

## 1. Introduction

A kombucha is fermented sugared tea with antimicrobial, antioxidant and probiotic properties [1, 2, 3, 4, 5, 6, 7, 8, 9, 10, 11]. The tea is fermented by a symbiotic community of bacteria and yeasts. The species of bacteria identified in kombucha include *Acetobacter xylinum, A. pasteurianus, A. acetic, A. intermedius, A. nitrogenifigens, Gluconobacter oxydans, G. kombucha, Leuconostoc* spp., *Allobaculum* spp., *Ruminococcaceae* spp., *Enterococcus* spp., *Propionibacterium* spp., *Bifidobacterium* spp., *Thermus* spp., see details in [12, 3]. Yeast species are *Zygossacharomyces* spp., *Brettanomyces* spp., *Saccharomyces* spp., *Sacchromycodes* spp., *Pichia* spp., *Candida* spp., *Schizosaccharomyces* spp., *Mycoderma* spp. [3, 13, 12, 14, 15]. The fermented liquid where zooglea lives include acetic acid, gluconic acid, lactic acid, vitamins B1, B2, B6, B12, C, ethanol, proteins, poliphenols, Cu, Fe, MN, Ni, Zn, and a range of anions, including F^-^, Cl^-^, Br^-^, I^-^, NO_3_^-^, HPO_3_^-^, SO_4_^-^ [16, 3, 17, 18]. These symbiotic community of bacteria and yeasts produces the cellulosic gelatinous mat, also known as biofilm, commensal biomass, tea-fungus, scoby and zooglea. We decided to study electrical activity of the zoogleal mats for the following reasons.

First, the zooglea mats has attracted interest from sustainable fashion and textile industry, because the mats’ properties are similar to properties of clothing materials [19, 20, 21, 22, 23, 24]. Some future applications of the mats might require some loci of a mat to remain alive for sensing purposes, where chemical, tactile or optical stimuli will be mapped onto patterns of electrical activity. Thus kombucha mats based wearables will be capable of detecting and signalling of changes in ambient air and light.

Second, recent discoveries and designs of living electronic devices [25, 26, 27] demonstrated that one can make a wide range of electronic devices with slime mould [28], fungi [29], plants [30], algae [31]. If kombucha zoogleal mats acted as ‘continuous’ sheets with non-linear electrical properties and endogenous electrical activity the mats would be fertile substrate for future living electronics and computing devices. Pioneer experiments with acetobacter colonies [32, 33] successfully demonstrated that bacterial films can be used for sensing purposes.

Third, kombucha zoogleal mats are unique eco-systems where yeasts and bacteria cohabit, compete and cooperate, and, as discussed in [12], the substrate can be used as a model of many species cooperation. In this sense, a kombucha mat can be seen as a single organism which coordinates and integrate all its parts into a coherent metabolic activities. Bio-electrical activity [34, 35, 36, 37] is proved to act as a vehicle of organisms’ integration, regeneration and proto-cognition. Thus, by studying electrical activity of kombucha mats we can uncover mechanisms of long-scale communication and, possibly, sensorial fusion and information processing in microbial populations comprised of a very large number of species. Electrical activity of bacterial film was reported as early as 1964 in relation to electrical currents generated by microbial films of dental plague [38]. Further studies demonstrated that ion channels conduct long-range electrical signals within bacterial biofilms via travelling waves of potassium [39]. Spatially extended electrical activity of kombucha mats might be an indicator of complex multi-species interactions.

The paper is structured as follows. We introduce experimental setup in Sect. 2. We analyse endogenous electrical activity of kombucha mats and the dynamic of the activity in response to chemical, optical and electrical stimulation in Sect. 3. In Sect. 4 we discuss the findings and outline pathways for further studies.

## 2. Methods

The kombucha zooglea was initially obtained from Freshly Fermented Ltd (Lee-on-the-Solent, PO13 9FU, UK) and then produced *in situ*. The infusion was prepared in the following proportions: 2% of tea (PG Tips UK), 5% sugar (Silver Spoon, PE2 9AY, UK), 1 L of boiled tap water. Containers with kombucha were kept at 20-23°C in darkness. Mother solution was changed every 7 days. Electrical activity of the kombucha was recorded using pairs of iridium-coated stainless steel sub-dermal needle electrodes (Spes Medica S.r.l., Italy), with twisted cables and ADC-24 (Pico Technology, UK) high-resolution data logger with a 24-bit analog-to-digital converter (Fig. 1a). We recorded electrical activity at one sample per second. During the recording, the logger has been doing as many measurements as possible (typically up to 600 per second) and saving the average value. We set the acquisition voltage range to 156 mV with an offset accuracy of 9 *µ*V at 1 Hz to maintain a gain error of 0.1%. Each electrode pair was considered independent with the noise-free resolution of 17 bits and conversion time of 60 ms. Each pair of electrodes, called channels, reported a difference of the electrical potential between the electrodes. Distance between electrodes was 1-2 cm.

**Figure 1:**
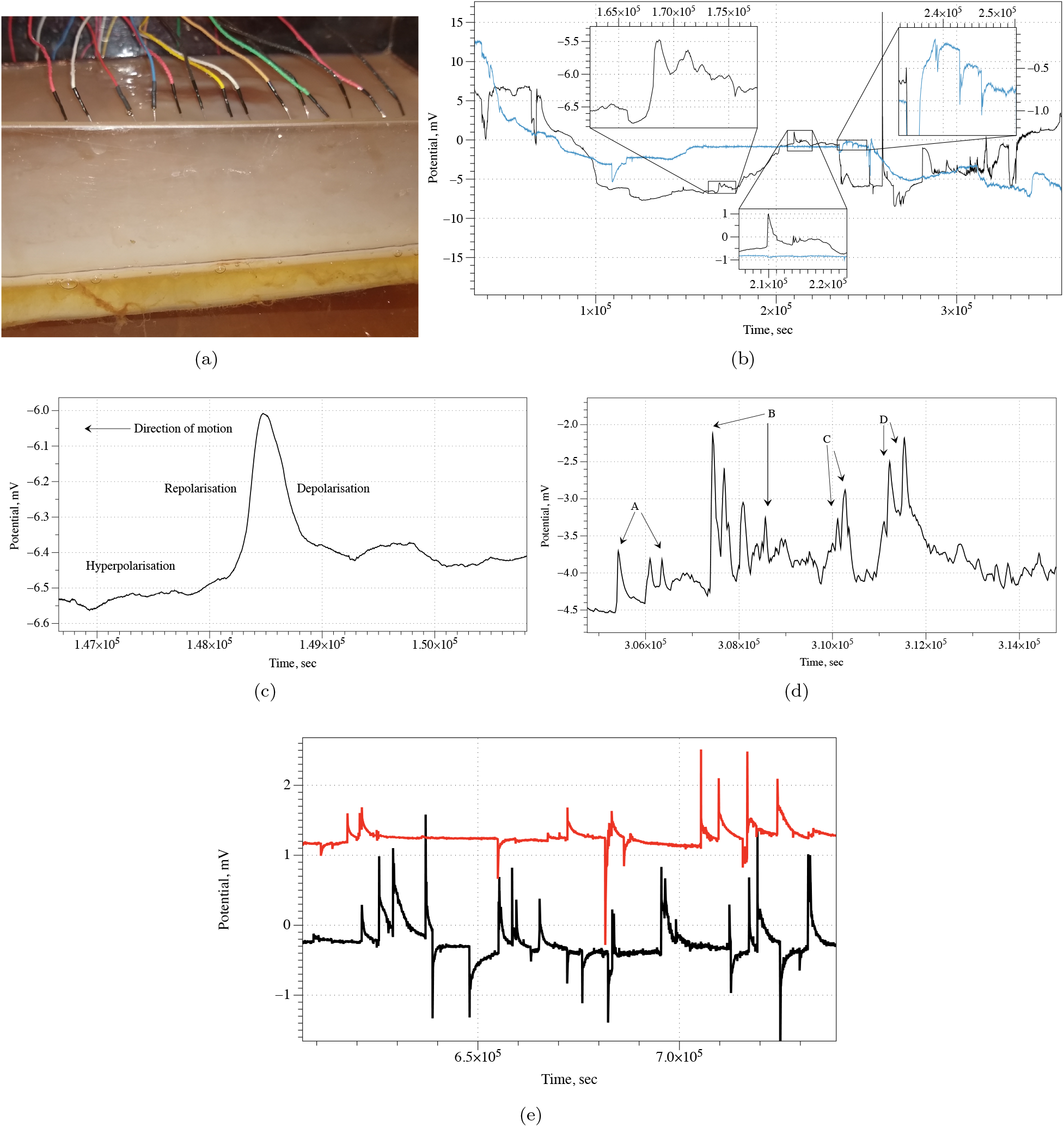
(a) Experimental setup: pairs of differential electrodes are inserted into kombucha mat. (b) Examples of recordings on two pairs of differential electrodes. (c) Action potential like spike of electrical potential recorded on kombucha mat. (d) Example of a group of spike trains. (e) Electrical activity recorded on a two pair of neighbouring differential electrodes.

For optical stimulation we used an aquarium light, array of 36 white LEDs, illumination on the fungal skin was 0.3 LUX. For chemical stimulation we used sucrose (Sigma Aldrich). Electrical stimulation was provided by PSD 30/3B high performance regulated DC power supply (BST Caltek Industrial Ltd.).

We have conducted 12 experiments, each experiment lasted at least seven days. In each experiment we recorded electrical activity of the fungi via four pair of differential electrodes, i.e. 48 seven days, or more, long recordings in total.

## 3. Results

### 3.1. Endogeneous spiking

We found that pairs of differential electrodes inserted in kombucha zoogleal mats show a rich dynamics of electrical activity. A slow dynamics is characterised by changes of electrical potential by 5-20 mV occurred over several days Fig. 1b. A fast dynamic is represented by spikes of electrical potential which duration ranges from 5 min to several hours. An example of the action-potential like spike with 0.5 mV, is shown in Fig. 1c, characteristic depolarisation (c. 12 min), repolarisation (c. 10 min) and repolarisation phase (c. 19 mins). The spikes often found grouped in trains of spikes. Examples of spike trains are show in inserts in Fig. 1a: a train of four spikes, top left insert, three pikes, top right insert, and two spikes, bottom insert, and Fig. 1d.

Parameters of the electrical spiking are as follows. Plot of spikes duration versus spikes amplitude in Fig. 2a shows that there is no apparent correlation between these two characteristics. Pearson correlation coefficient R is 0.149, whilst a correlation is positive, the relationship between spike amplitude and width is weak. A distribution of spike amplitudes is shows in Fig. 2b. Average amplitude is 0.8 mV, median 0.4 mV, *σ* = 0.6. The amplitude distribution is far from normal: the value of the K-S test statistic *D* is 0.2, *p* < 0.03. Average duration of a spike is 9.8 min, median 3.6 min, *σ* = 10.5 min. The spike amplitude distribution is highly asymmetric, skewness is 0.7, with long tail, kurtosis is −1.025572. The spike duration distribution is also not normal, *D* = 0.35, *p* < .00001, and asymmetric: skewness is 1.9 and kurtosis 3.7.

**Figure 2:**
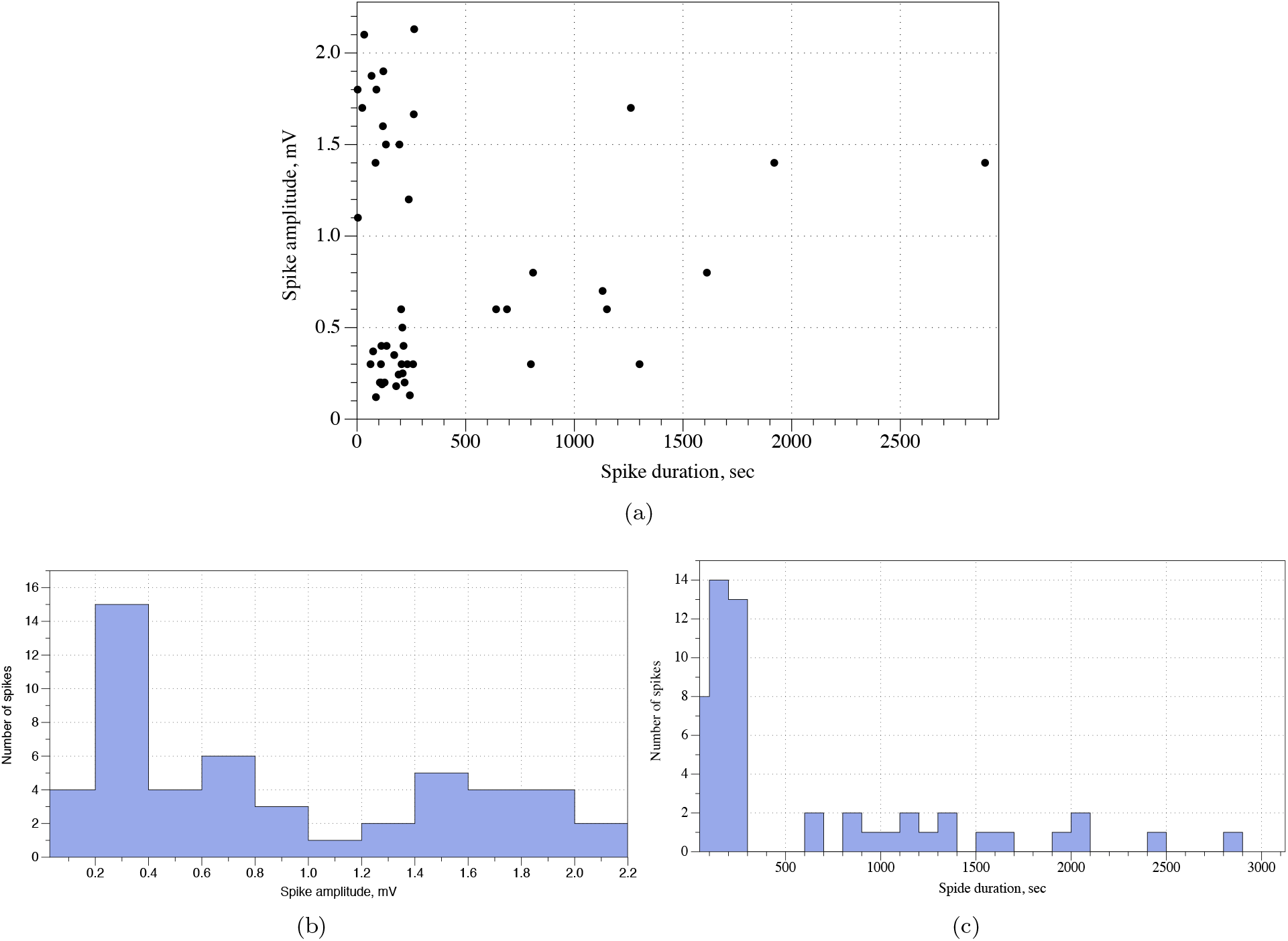
Characteristics of electrical spiking behaviour. (a) Spike duration versus amplitude. (b) Distribution of spike amplitudes. (c) Distribution of spike durations.

There are 3.2 spikes in average in a train of spikes, *σ* = 1.3, median 3. In more details, the set of spike trains consists of two distinctive subsets: fast trains (60%) and slow trains (30%). In fast trains, an average spike width is 2.8 min, median is 3.2 min, *σ* = 1.1 min. In slow trains, an average spike width is 24.2 min, median is 22.6 min, *σ* = 7.6 min. For all spike trains studied distance *d* between adjacent spikes is a function of spikes width *w*: *d* = *a* ∗ *w*, where average *a* is 1.1, median 1.1 and *σ* = 0.3.

Does the electrical activity propagates along zoogleal mats? Some recordings, e.g. Fig. 1e, demonstrate that out of 16 spikes recorded on one pair of differential electrodes (red plot in Fig. 1e) 13 spikes are mirrored, with delay, at the nearby pair of differential electrodes (black plot). Average delay is 1.2 hr, median 1.1 hr, *σ* = 0.7. Assuming a distance between two neighbouring pairs of differential electrodes does not exceed 3 cm, we can propose that electrical activity propagates with the speed of 2.4 cm/hr.

### 3.2. Response to stimulation

Let us consider a typical response to stimulation with sucrose (Fig. 3a). In the intact state the zoogleal mat exhibits low amplitude 0.2-0.8 mV electrical activity. In c. 23 hr after application of 20 g glucose, the mat started to show high amplitude oscillations. Average duration of a spike is 5.3 hr (*σ* = 1.7), average amplitude of a spike is 4 mV (*σ* = 2.7), and average distance between the spikes is 5.7 hr (*σ* = 3). These high amplitude low frequency electrical responses ended in c. 5-6 days after the stimulation.

**Figure 3:**
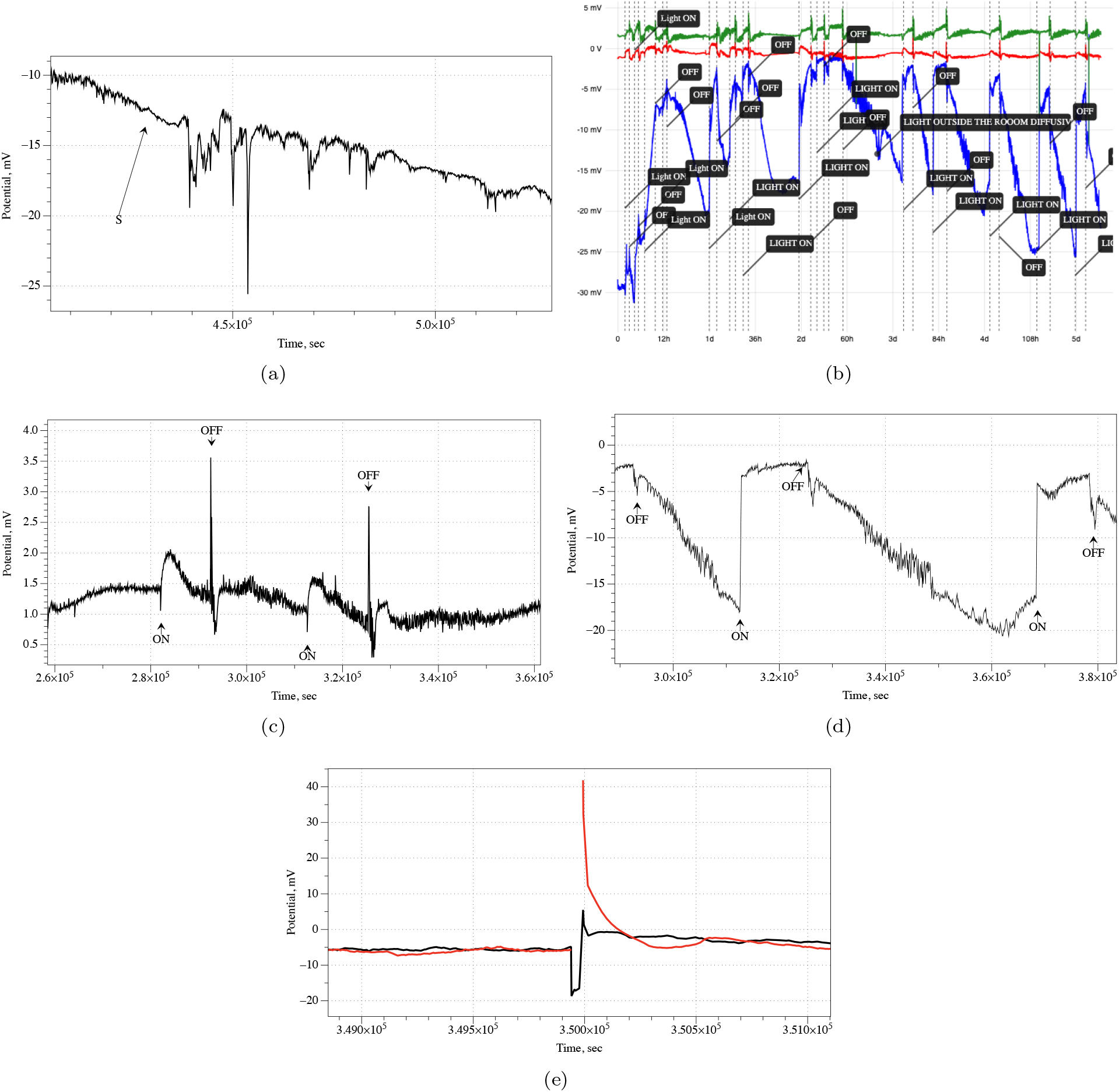
Responses to stimulation. (a) Application of sucrose. (b) Stimulation with light, screenshot of PicoLog 6 running running electrical recording, moments when light was on and off are indicated on the plot. Channel closet to the source of light is blue, farthest is green, and median is red. (c) Electrical response to illumination on the differential electrodes pairs being at lest 6 cm away from the source of light. (d) Electrical response to illumination on the differential electrodes pair being directly under the source of light. (e) Response to stimulation with 20 VDC, 20 sec.

A response to optical stimulation, recorded on three pairs of differential electrodes is shown in Fig. 3b. The recordings Fig. 3b and zoomed parts of the activity several centimetres away from the source of illumination (Fig. 3c) and directly under the source of illumination (Fig. 3c) give us an indication that an intensity of the response might be dependent on a distance from the source of light. At the site under the source of illumination the potential raise by 15.3 mV in average, *σ* = 1.9 in 0.7 hr in average, *σ* = 0.6. At other site of the zoogleal mat, the potential raises by 0.9 mV in average, *σ* = 0.4 in 3 hr in average, *σ* = 0.3. When the light is switched the potential at the loci being directly under the source of stimulation drops by 15.6 mV, *σ* = 4 in 7 hr, *σ* = 2.7. The drop of the potential at other site is 1.3 mV in average, *σ* = 0.2, in 0.4 h, *σ* = 0.1.

We did not observe any changes in spiking activity of kombucha mats in response to electrical stimulation, 20 VDC c. 20 sec. Typical responses, on two pairs of differential electrodes, are shown in Fig. 3e. Whilst no spiking responses have been detected, it is evident that zoogleal mats exhibit properties of capacitors. Stimulation with 20 VDC cause the potential to raise by 23.4 mV in average, *σ* = 17.2. The potential drops gradually, with an average discharge duration of 3.9 hr, *σ* = 0.1.

## 4. Discussion

We recorded and analysed action-potential like spikes of electrical activity in kombucha zoogleal mats. The spikes of electrical activity are in fact two-dimensional excitation waves propagating in kombucha mats recorded on differential electrodes as spike. This is illustrated in Fig. 4. The excitation waves are numerically modelled using FitzHugh-Nagumo equation, see details of the model in [40, 41]. A pair of differential electrodes is shown in Fig. 4a. The pair records a potential difference between right and left electrode. Thus, when an excitation wave propagates from the left to the right (Fig. 4b) we see a drop of the potential followed by a raise of the potential and then one more spike is manifested in a raise of the potential followed by a fall of the potential (dotted plot in Fig. 4d). When the excitation wave propagates from the right to the left (Fig. 4c) the spikes appear in a the reverse order of configurations (solid plot in Fig. 4d). Exact geometries of the spikes depend on the angle the excitation waves cross the pair of differential electrodes. This is illustrated in Fig. 4efg. Distance between electrodes affects amplitudes of spikes recorded, as seen by comparing plots Fig. 4f and Fig. 4g. Several waves colliding at the loci of differential electrode pairs might produce no spikes due to equal potential on both electrodes.

**Figure 4:**
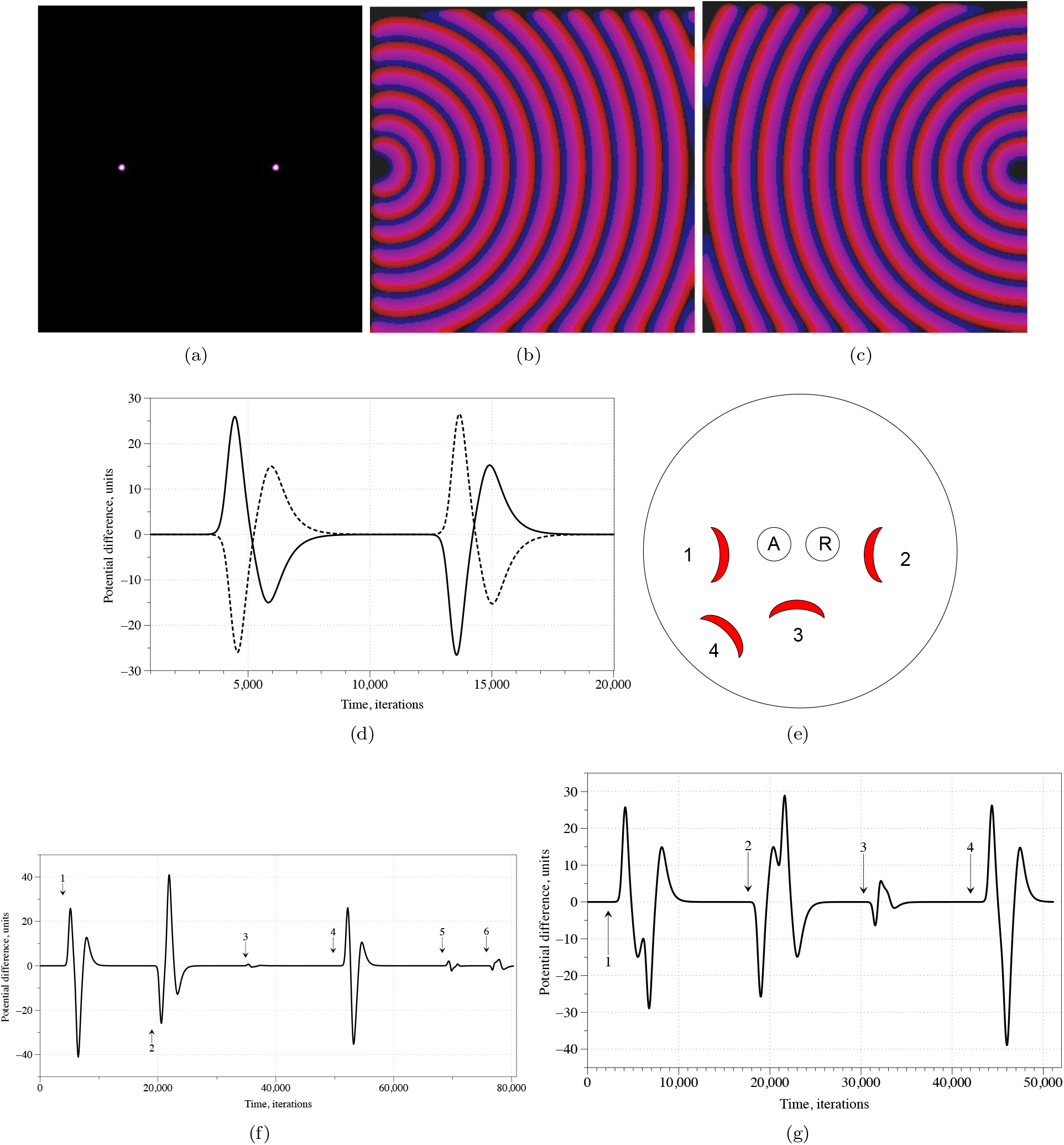
(abc) An illustration on the nature of electrical potential spikes. Distance between electrodes is 140 nodes. (a) Position of electrodes. (b) Time lapse of the wave propagating from the left to the right. (c) Time lapse of the wave propagating from the right to the left. (d) Potential difference recorded on the electrodes for both positions of the excitation source. Solid line is a potential recorded when wave propagates from the right to the left, see video in https://youtube.com/shorts/J1tVYSEKvQo? feature=share; dotted line – frome the left to the right, see video in https://youtube.com/shorts/n0xA_KnimeQ?feature=share. (e) A scheme of excitation sources positions and a pair of differential electrodes. (fg) Electrical potential difference recorded on the electrodes, distance between electrodes is 20 nodes (f), 40 nodes (g). Spikes specific to only location of excitation source are labelled by numbers corresponding to the excitation sources. Spikes 5 and 6 correspond to the situation when several wave fronts from different directions collide near the vicinity of the electrodes. See an example here https://youtube.com/shorts/b-xTeXR5R8U?feature=share.

Biophysical origins of excitation waves in kombucha mats might be as following.

First, as proposed in [39], the waves of depolarisation, emerging due to metabolically triggered release of potassium, travelling in a bacterial film coordinate the cells’ metabolic states. The excitation wave, recorded as spikes, have been also observed in acetobacter colonies [32, 33].

Second, several species of yeasts are found in kombucha mats. Glicolytic oscillations of yeasts are reflected in oscillations of their resistance and capacitance [42]. Most likely there is also some electrical potential difference generated during the glycolysis. Our personal unpublished studies found oscillations of electrical potential of kefir grains (Fig. 5a), where duration of spikes varies from 10 to 30 hours, amplitude from 0.5 to 1.5mV. Such periods of spikes match duration of kombucha mat spiking when subjected to stimulation with sucrose (Subsect. 3.2). Kombucha mats often exhibit bursting mode electrical activity (Fig. 5b) which resembles bursting modes of neurons, as illustrated in the model of Izhikevich neuron [43] in Fig. 5c. A model presented in [44] indicates that a feedback of the cellulose matrix Ca^2+^ onto ion channels coupled with slow oscillations driven by glycolysis can lead to such slow bursting modes.

**Figure 5:**
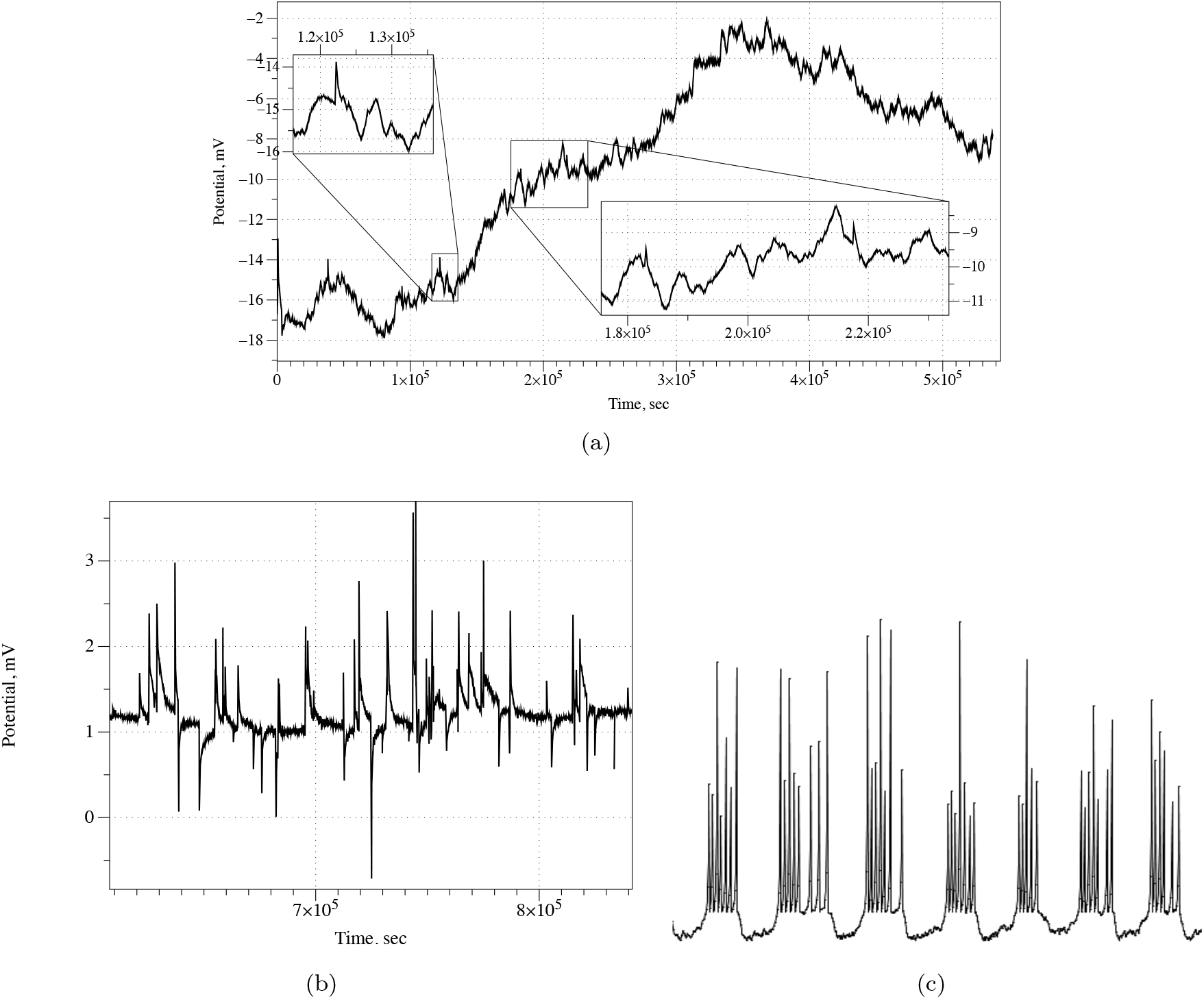
(a) Oscillations in Kefir grains. (b) Bursting mode of electrical activity of kombucha mat. (c) Bursting in Izhikevich neuron.

Third, some evidence is provided in [45], where formation, propagation and on-collision annihilation of NADH and proton waves have demonstrated in spatially distributed non-mixed yeast culture. Proton waves in yeast culture propagate with the speed 5*·*10^−4^ cm/sec [45]. At the end of the Sec. 3.1 we proposed that excitation waves propagate in kombucha mat with the speed c. 2.4 cm/h, i.e. c. 7*·*10^−3^ cm/sec. The proton waves in yeast culture propagate an order faster [45] however such observation have been made at a microscopic scale. Presence of cellulose matrix in kombucha mats can substantially slow down propagation of the excitation waves. Also, it worth to note that estimated [46] electrical activity wave in fungi is 5*·*10^−2^ cm/sec, which is several orders slower than speed of a typical action potential in plants: from 0.005 m/sec to 0.2 m/sec [47], but just an order faster than speed of electrical activity propagation in kombucha mats.

What would be directions of future developments related to electrical activity of zoogleal kombucha mats? The kombucha mats are very convenient for experimentation and, in principle, do not require any sterile environment and can be experimented with in literally any environment, including electronics and engineering laboratory. This is because the kombucha shows antimicrobial activity against *H. pylori, S. typhimurium, S. aureus, A. tumefaciens, B. cereus, S. sonnei, S. enteritidis, E. coli* [48]. Thus one could use living kombucha mats in developing living bioelectronic devices, substrate with active and non-linear electrical properties. In future, we will explore sensing and computing properties of kombucha mats. To fully evaluate sensing abilities of the mat, we will aim to reconstruct a one-to-one mapping between key chemical, physical and optical stimuli and patterns of electrical activity of the mats.

Computational properties of kombucha zoogleal mats could be evaluated by using three approaches. First, we can applying our developed techniques for mining logical functions [49, 50] to construct distributions of many-input Boolean functions. Second, we can prototype neuromorphic circuits with kombucha mats. There is a high likelihood of kombucha mats exhibiting memristive properties (similar to memristive properties of slime mould [51] and fungi [52]). The memristive properties would further imply [53, 54, 55, 56] that it is possible to implement learning, memory and construct synaptic connections in the zoogleal mats. Third, we can adopt reaction-diffusion algorithms [57] to implement spatial computing, e.g. some tasks of computational geometry and image processing, with kombucha. As demonstrated in [45] proton waves in yeast culture travel and interact alike oxidation waves in Belousov-Zhabotinsky (BZ) medium. Therefore, architectures of most experimental prototypes of BZ medium based computers [58, 59, 60, 61] can be realised with kombucha zoogleal mats.

